# Prognostic significances of interleukin-17-producing cells and Th17 cells in malignant cancers: a meta-analysis of the literatures

**DOI:** 10.1101/869776

**Authors:** Yong Luo, Ting Yu, Cheng Yi, Huashan Shi

**Affiliations:** State Key Laboratory of Biotherapy and Department of Head and Neck Oncology, West China Hospital, West China School of Medicine, Sichuan University, Chengdu, Sichuan, China; Department of Pathology and Laboratory of Pathology, State Key Laboratory of Biotherapy, West China Hospital, West China School of Medicine, Sichuan University, Chengdu, Sichuan, China; Department of Biotherapy, Cancer Center, West China Hospital, Sichuan University, Chengdu, China; Division of Hematology and Oncology, Mayo Clinic, Jacksonville, USA

## Abstract

**Background and purpose:** As a proinflammatory factor, interleukin-17 (IL-17) can play a role in both tumor promotion and suppression. IL-17 is traditionally regarded as secreting mainly by CD4^+^ T helper cells (Th17 cells), while other immune subsets have been proved to produce IL-17, called IL-17^+^ cells. Considerable studies have drawn controversial conclusions about association between IL-17^+^/Th17 cells and prognosis of cancer patients. This meta-analysis was performed to systematically and quantitatively analyze prognostic values of IL-17^+^ cells and Th17 cells in cancer patients.

**Methods:** A comprehensive retrieval was conducted in Pubmed (MEDLINE) and EMBASE databases. Pooled risk ratios (RRs) or hazard ratios (HRs) and corresponding 95% confidence intervals (CI) were calculated to evaluate the prognostic values of IL-17^+^ cells and Th17 cells in cancer patients.

**Results:** A total of 42 studies with 5039 patitents were included. High IL-17^+^ cells was significantly associated with tumor recurrence (RR = 4.23, 95% CI [1.58, 11.35]), worse disease free survival (DFS) (HR = 1.84, 95% CI [1.22, 2.77]) and overall survival (OS) (HR = 1.39, 95% CI [1.04, 1.87]), especially in cancers of digestive system. Besides, no significant difference was observed between high IL-17^+^ cells and histological grade, lymph node metastasis, tumor volume, clinical stages or distant metastasis. Moreover, there was no significant difference in OS between high and low Th17 cells in cancer patients (HR = 0.93, 95% CI [0.58, 1.49]).

**Conclusions:** This meta-analysis suggests high IL-17^+^ cells could be an indicator for worse survival in patients with malignant cancers, especially with cancers of digestive system. Although high Th17 cells appears to have non-statistically significance on prognosis, more clinical studies should be implemented to investigate the underlying function of Th17 cells within tumor microenvironment. This study put forward a new insight for potential application of anti-IL-17 target therapy in cancer therapeutics.

## Introduction

With the prolongation of longevity, cancer has gradually been the leading cause of death and major burden of public health system worldwide [1]. The interactions between malignant cells and non-malignant cells in tumor mass constitute tumor microenvironment, which is highly important for tumorigenesis, cancer progression and responses to treatments [2]. Multiple immune cells and relevant cytokines form the essential component of tumor microenvironment. Despite of their crucial role in anti-tumor responses, these immune cells involving lymphocytes, macrophages and neutrophils can also possess a dynamic and tumor-promoting capacity almost in all solid tumors [3,4]. The success of immunotherapy in cancer patients has attracted enormous attentions to tumor immune microenvironment, aiming at finding more effective therapy and novel therapeutic targets.

Interleukin 17 (IL-17), also named cytotoxic T lymphocyte-associated-8 (CTLA-8), was firstly recognized in 1993 [5]. The IL-17 family of cytokines contains six membranes: IL-17A-F, of which IL-17 was recently renamed as IL-17A [6]. Among the IL-17 family membranes, only IL-17F and IL-17A share some homology and overlapping functions. Worked as a proinflammatory factor, IL-17 can attract neutrophils and stimulate inflammatory responses [7]. In the last decade, IL-17 became striking with the finding of a novel population of IL-17-producing CD4^+^ T helper cells, which termed as type 17 T helper (Th17) cells. Th17 cells can secrete IL-17A, IL-17F and IL-22 to induce inflammatory response to against pathogens that could not be handled properly by Th1 and Th2 cells [8]. Th17 cells are generally regarded as the major producer of IL-17, while other immune subsets such as IL-17-producing CD8^+^ T cells (Tc17 cells), natural killer (NK) cells, γδTcells, and innate lymphoid cells (ILC) also have been proved to produce IL-17 [9,10].

Although the proinflammatory character of IL-17 is crucial for host-protection, the excess IL-17 secreting has been shown to be connected with autoimmune disease and cancer development [11,12]. Antibodies against IL-17A, such as secukinumab or ixekizumab have been approved for treating moderate to severe psoriasis [13,14]. Chronic inflammatory conditions are also tightly associated with high risks of cancer progression [15]. In recently years, a growing body of studies has shown IL-17-producting cells (IL-17^+^ cells) are extensively existed in various inflammation-related cancers, such as lung cancer, hepatocellular carcinoma (HCC), colorectal cancer (CRC) and breast cancer [16–19]. More infiltration of IL-17^+^ cells was founded in intratumor tissues than that in peritumor nontumor tissues [20,21]. Besides, the level of Th17 cells was also founded higher in peripheral blood of cancer patients than that in normal control patients [18]. IL-17^+^ cells seem to produce a dual role in tumor. On the one hand, the main function of IL-17 in tumor microenvironment tends to promote tumor progression [22]. The pro-tumor function of IL-17 mainly depends on its proangiogenic capability which mediated by surrounding fibroblasts and endothelial cells. IL-17 can induce the production of vascular endothelial growth factor (VEGF) by fibroblasts [23,24]. The IL-17-VEGF loop not only facilitates the fibroblast-induced angiogenesis, but also induces other angiogenic factors, such as TGF-β, IL-6 and PGE_2_ [25,26]. The TGF-βin turn improves the VEGF receptivity via up-regulating expression of VEGF receptor [27]. Significant positive relationships have been discovered between microvessels density and IL-17^+^ cells density within tumor tissues [28]. IL-17 has been reported to promote breast cancer progression by inducing the recruitment of tumorigenic neutrophils [29]. On the other hand, IL-17 also generates some tumor-inhibitory abilities. IL-17 is able to stimulate the production of IL-12 from macrophages, which is associated with activating tumor-specific cytotoxic T lymphocytes (CTLs) [30,31]. Besides, IL-17 also accelerates the dendritic cells (DCs) maturation by up-regulating costimulatory molecules and MHC class II Ags [32].

Meanwhile, some clinical evidences also indicate the controversial prognostic role of IL-17^+^ cells and Th17 cells in cancer patients. High levels of Th17 cells in peripheral blood or high density of intratumoral IL-17^+^ cells has been reported to be associated with worse survival in patients with HCC [19], while better survival also has been observed in cancer patients with high level of IL-17^+^ cells or Th17 cells [18,33]. Furthermore, some researchers have already noted that IL-17 and Th17 cells are not synonymous [34,35]. Therefore, this meta-analysis was conducted to systematically study the significances of IL-17^+^ cells and Th17 cells levels in clinicopathological features and survival of cancer patients, as well as further understanding of the differences between IL-17^+^ cells and Th17 cells in prognostic values.

## Materials and methods

### Search strategy

We conducted a comprehensive literature retrieve in the Pubmed (MEDLINE) and EMBASE databases, basing on the terms “T helper 17 (Th17) cells [Title/Abstract]”, “Interleukin-17 (IL-17) [Title/Abstract]”, “tumor/cancer/neoplasm/carcinoma [Title/Abstract]” without any limitations of language and publication date. The most recently retrieved literatures were updated on October 30^th^, 2019. The references from all electable studies were manually screened in case of missing any potentially included studies. In this study, we also referred the PRISMA and EQUATOR Reporting guidelines in order to follow the criterions of meta-analysis.

### Criteria for inclusion and exclusion

In this meta-analysis, include studies need to satisfy the following criteria: (1) included patients were pathologically diagnosed with malignant cancers; (2) expression of IL-17^+^ cells or Th17 cells was assessed in cancer patients; (3) the association between expression of IL-17^+^/Th17 cells and clinicopathological features or survival data were analyzed. Studies were excluded if meeting one of the following criteria: (1) study was published in the form of review, letter, conference abstract, editorial or case report; (2) data between expression of IL-17^+^/Th17 cells and clinicopathological features or survival data were unable to obtain; (3) the duplicated studies or studies with overlapping patients in which the one with smaller sample size should be excluded.

### Data extraction

Data were independently extracted by two reviewers, complying with a standard procedure of data collection. The following items were gathered: author, publication year, country, study design, patients number, sample source, methods used for measuring IL-17^+^ cells or Th17 cells expression, cut-off values, median follow-up time, related data of clinicopathological features and survival. The unconformities between the data extracted by two reviewers were resolved by serious discussion. Furthermore, the qualities of included studies were scored according to the Newcastle-Ottawa quality assessment scale (NOS scale), and defined the high quality was scored more than 5 points.

### Statistical analysis

In this study, a total of nine groups were analyzed for assessing the prognostic values of IL-17^+^ cells and Th17 cells in malignant cancers, involving histological grade, lymph node status, tumor volume (T stages), clinical stages, distant metastasis, tumor recurrence, disease free survival (DFS) and overall survival (OS). For quantitatively aggregating these results, the impacts of IL-17^+^ cells level on clinicalpathological features were assessed by using risk ratio (RR), and the influences of IL-17^+^ cells or Th17 cells level on patients survival were analyzed by using hazard ratio (HR), with their 95% confidence interval (CI). The values of RRs/HRs and 95% CI were directly extracted from the reported data in include studies. If those data were unable to obtain, the total number of events and number of patient at risk in each group were collected for calculating the RRs and their 95% CIs, according to the methods described before [36]. Moreover, HRs and 95% CIs of survival data were calculated by using Engauge Digitizer (free software downloaded from http://sourceforge.net) to read the Kaplan-Meier curves, when the provided data were only represented in graphical forms. The pooled RRs/HRs and theirs 95% CIs were shown as forest plots. If the pooled RR/HR was > 1, it indicated a worse prognosis/survival for patients with high IL-17^+^ cells or Th17 cells. If the 95% CIs of RRs or HRs did not overlap 1, the impacts of high IL-17^+^ cells or Th17 cells on clinicopathological features or survival were identified as statistically significant. Furthermore, the *I*^2^ metric and Q statistic were applied for evaluating the heterogeneities. A fixed effect model was employed if the heterogeneity was unobvious (*I*^2^ < 50% or P >0.1), or a random effect model was chosen (*I*^2^ > 50% or P <0.1). Besides, the publication bias of each group was also analyzed via the funnel plots (Begg’s test) and Egger’s linear regression method (Egger’s test), of which a p vale of <0.05 was thought to be statistically significant [37]. The software of STATA version 12.0 was used for this meta-analysis.

## Results

### Study selection and characteristic analysis

The primary search strategy retrieved 12474 studies, and 743 studies were carefully viewed in full text. Finally, a total of 42 studies were included in this meta-analysis. The Figure 1 showed the whole flow chart of literature screening. The total number of included patients was 5039, ranging from 25 to 458 patients per study. The general characteristics and NOS scales of the included studies about IL-17^+^ cells (Table 1) and Th17 cells (Table 2) were summarized in tables. Of the included studies, most of the studies were prospective cohort studies (n = 30), while 12 studies were retrospective studies. The majority of patients in the eligible studies had not received any chemoradiotherapy, immunotherapy or glucocorticoids before the sampling from blood or tumor tissues. Overall, a total of 34 studies investigated the prognostic value of IL17^+^ cells in clinicopathological features and survival of cancer patients, and 10 studies investigated the prognostic values of Th17 cells in survival, with 2 studies discussing both IL-17^+^ cells and Th17 cells. As for NOS scale, all of the included studies scored more than 5 points, which were considered as qualified enough for this meta-analysis. Table 3 summarized all of clinical outcomes analyzed in this study.

**Fig 1.**
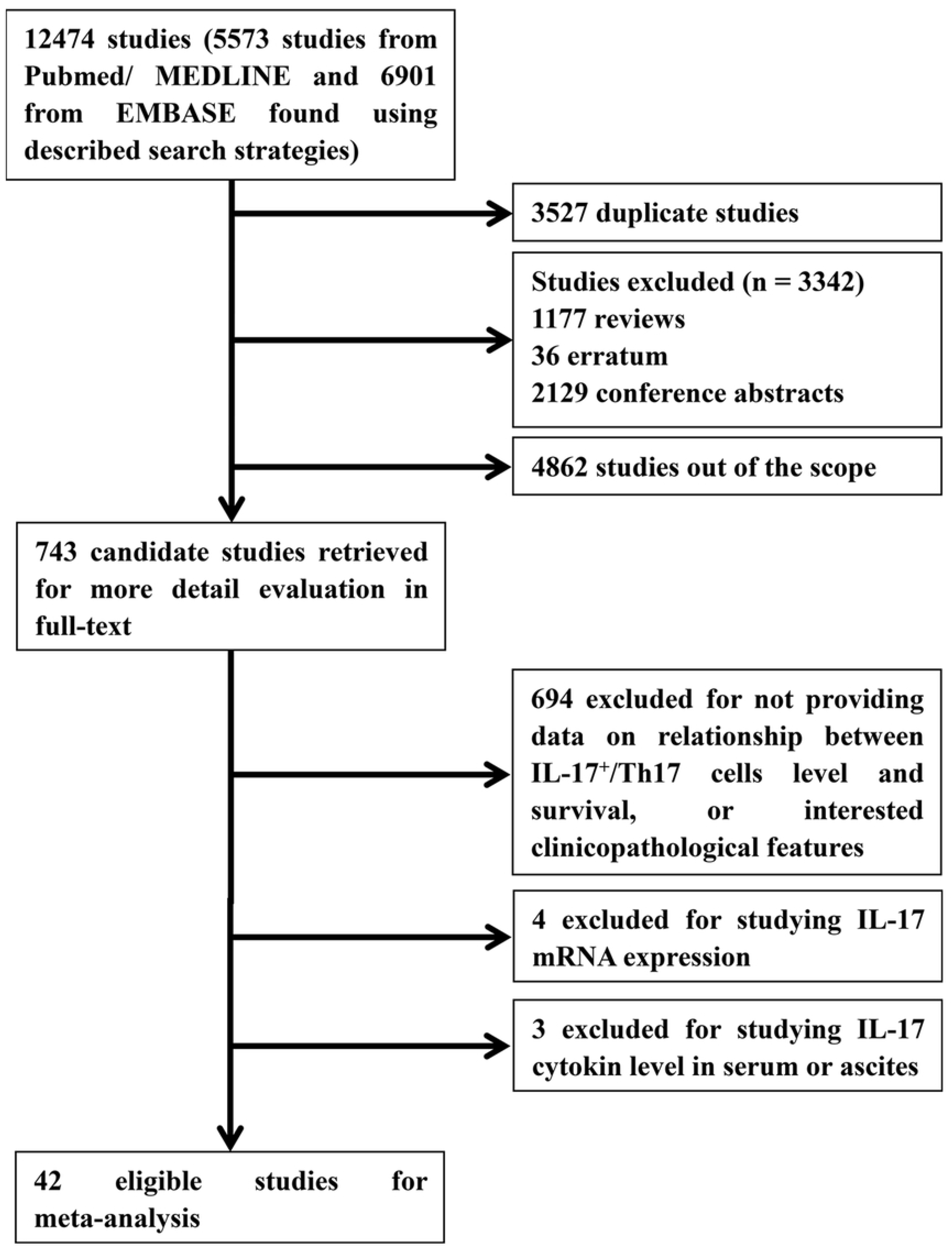
Flow chart of the literature retrieval and selection of included studies.

**Table 1.**
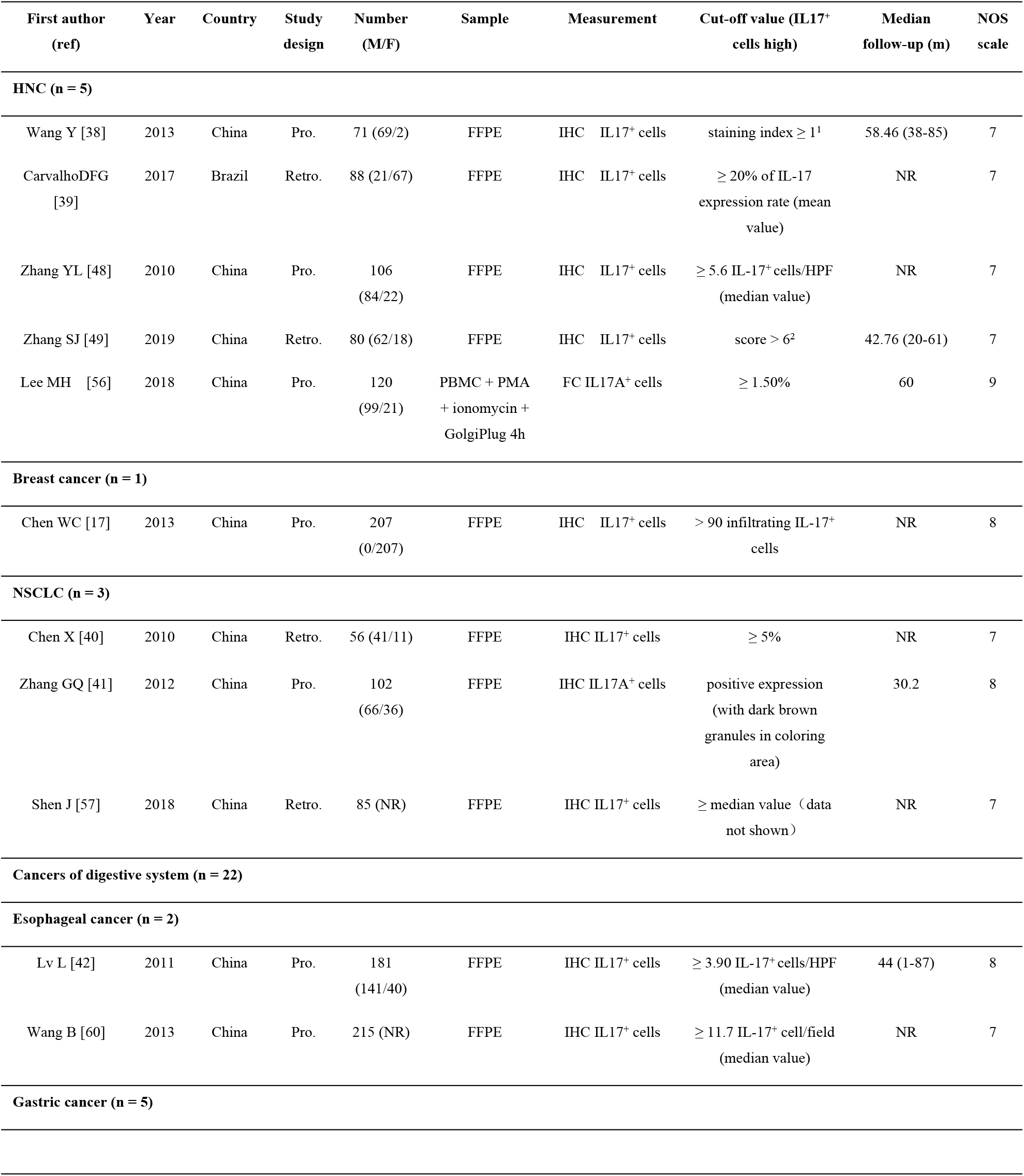
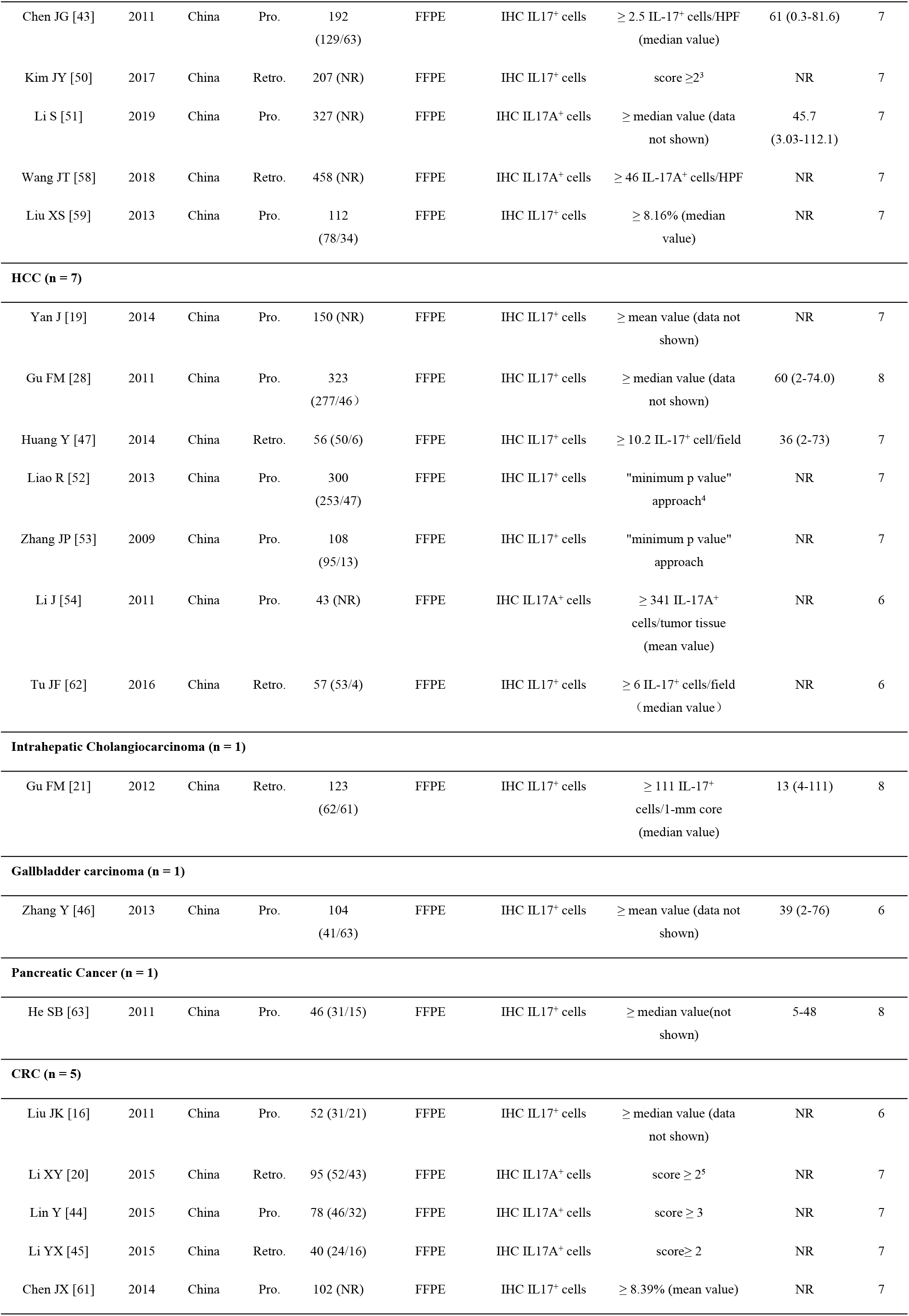
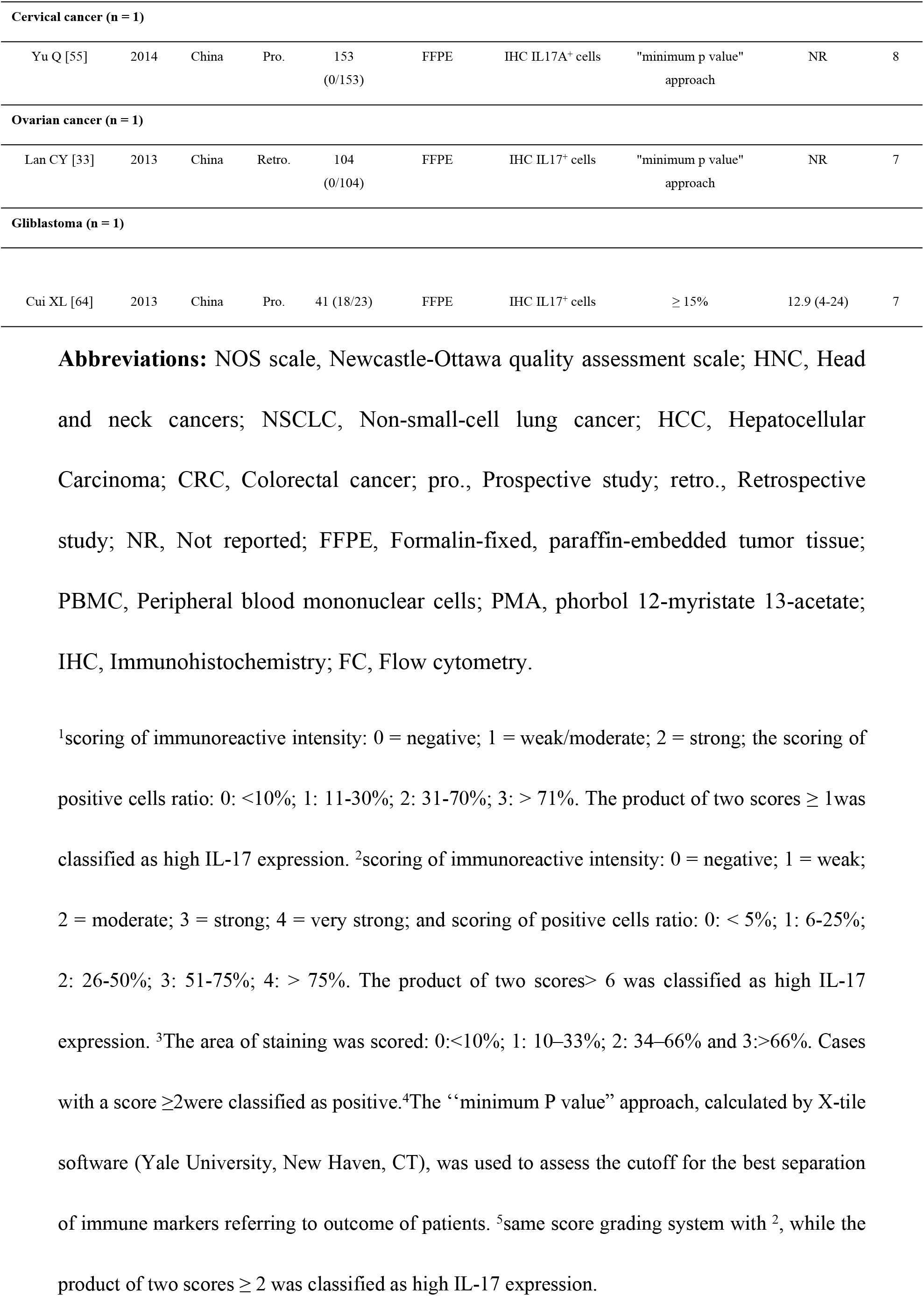
Main characteristics of the included studies about IL-17^+^ cells in this meta-analysis.

| First author (ref) | Year | Country | Study design | Number (M/F) | Sample | Measurement | Cut-off value (IL17<sup>+</sup> cells high) | Median follow-up (m) | NOS scale |
| --- | --- | --- | --- | --- | --- | --- | --- | --- | --- |
| **HNC (n = 5)** |  |  |  |  |  |  |  |  |  |
| Wang Y [38] | 2013 | China | Pro. | 71 (69/2) | FFPE | IHC IL17<sup>+</sup> cells | staining index $\geq 1^1$ | 58.46 (38-85) | 7 |
| CarvalhoDFG [39] | 2017 | Brazil | Retro. | 88 (21/67) | FFPE | IHC IL17<sup>+</sup> cells | $\geq 20\%$ of IL-17 expression rate (mean value) | NR | 7 |
| Zhang YL [48] | 2010 | China | Pro. | 106 (84/22) | FFPE | IHC IL17<sup>+</sup> cells | $\geq 5.6$ IL-17<sup>+</sup> cells/HPF (median value) | NR | 7 |
| Zhang SJ [49] | 2019 | China | Retro. | 80 (62/18) | FFPE | IHC IL17<sup>+</sup> cells | score $> 6^2$ | 42.76 (20-61) | 7 |
| Lee MH [56] | 2018 | China | Pro. | 120 (99/21) | PBMC + PMA + ionomycin + GolgiPlug 4h | FC IL17A<sup>+</sup> cells | $\geq 1.50\%$ | 60 | 9 |
| **Breast cancer (n = 1)** |  |  |  |  |  |  |  |  |  |
| Chen WC [17] | 2013 | China | Pro. | 207 (0/207) | FFPE | IHC IL17<sup>+</sup> cells | $> 90$ infiltrating IL-17<sup>+</sup> cells | NR | 8 |
| **NSCLC (n = 3)** |  |  |  |  |  |  |  |  |  |
| Chen X [40] | 2010 | China | Retro. | 56 (41/11) | FFPE | IHC IL17<sup>+</sup> cells | $\geq 5\%$ | NR | 7 |
| Zhang GQ [41] | 2012 | China | Pro. | 102 (66/36) | FFPE | IHC IL17A<sup>+</sup> cells | positive expression (with dark brown granules in coloring area) | 30.2 | 8 |
| Shen J [57] | 2018 | China | Retro. | 85 (NR) | FFPE | IHC IL17<sup>+</sup> cells | ≥ median value (data not shown) | NR | 7 |
| **Cancers of digestive system (n = 22)** |  |  |  |  |  |  |  |  |  |
| **Esophageal cancer (n = 2)** |  |  |  |  |  |  |  |  |  |
| Lv L [42] | 2011 | China | Pro. | 181 (141/40) | FFPE | IHC IL17<sup>+</sup> cells | $\geq 3.90$ IL-17<sup>+</sup> cells/HPF (median value) | 44 (1-87) | 8 |
| Wang B [60] | 2013 | China | Pro. | 215 (NR) | FFPE | IHC IL17<sup>+</sup> cells | $\geq 11.7$ IL-17<sup>+</sup> cell/field (median value) | NR | 7 |
| **Gastric cancer (n = 5)** |  |  |  |  |  |  |  |  |  |
| Chen JG [43] | 2011 | China | Pro. | 192<br>(129/63) | FFPE | IHC IL17<sup>+</sup> cells | ≥ 2.5 IL-17<sup>+</sup> cells/HPF<br>(median value) | 61 (0.3-81.6) | 7 |
| Kim JY [50] | 2017 | China | Retro. | 207 (NR) | FFPE | IHC IL17<sup>+</sup> cells | score ≥ 2<sup>3</sup> | NR | 7 |
| Li S [51] | 2019 | China | Pro. | 327 (NR) | FFPE | IHC IL17A<sup>+</sup> cells | ≥ median value (data<br>not shown) | 45.7<br>(3.03-112.1) | 7 |
| Wang JT [58] | 2018 | China | Retro. | 458 (NR) | FFPE | IHC IL17A<sup>+</sup> cells | ≥ 46 IL-17A<sup>+</sup> cells/HPF | NR | 7 |
| Liu XS [59] | 2013 | China | Pro. | 112<br>(78/34) | FFPE | IHC IL17<sup>+</sup> cells | ≥ 8.16% (median<br>value) | NR | 7 |
| **HCC (n = 7)** |  |  |  |  |  |  |  |  |  |
| Yan J [19] | 2014 | China | Pro. | 150 (NR) | FFPE | IHC IL17<sup>+</sup> cells | ≥ mean value (data not<br>shown) | NR | 7 |
| Gu FM [28] | 2011 | China | Pro. | 323<br>(277/46) | FFPE | IHC IL17<sup>+</sup> cells | ≥ median value (data<br>not shown) | 60 (2-74.0) | 8 |
| Huang Y [47] | 2014 | China | Retro. | 56 (50/6) | FFPE | IHC IL17<sup>+</sup> cells | ≥ 10.2 IL-17<sup>+</sup> cell/field | 36 (2-73) | 7 |
| Liao R [52] | 2013 | China | Pro. | 300<br>(253/47) | FFPE | IHC IL17<sup>+</sup> cells | "minimum p value"<br>approach<sup>†</sup> | NR | 7 |
| Zhang JP [53] | 2009 | China | Pro. | 108<br>(95/13) | FFPE | IHC IL17<sup>+</sup> cells | "minimum p value"<br>approach | NR | 7 |
| Li J [54] | 2011 | China | Pro. | 43 (NR) | FFPE | IHC IL17A<sup>+</sup> cells | ≥ 341 IL-17A<sup>+</sup><br>cells/tumor tissue<br>(mean value) | NR | 6 |
| Tu JF [62] | 2016 | China | Retro. | 57 (53/4) | FFPE | IHC IL17<sup>+</sup> cells | ≥ 6 IL-17<sup>+</sup> cells/field<br>(median value) | NR | 6 |
| **Intrahepatic Cholangiocarcinoma (n = 1)** |  |  |  |  |  |  |  |  |  |
| Gu FM [21] | 2012 | China | Retro. | 123<br>(62/61) | FFPE | IHC IL17<sup>+</sup> cells | ≥ 111 IL-17<sup>+</sup><br>cells/1-mm core<br>(median value) | 13 (4-111) | 8 |
| **Gallbladder carcinoma (n = 1)** |  |  |  |  |  |  |  |  |  |
| Zhang Y [46] | 2013 | China | Pro. | 104<br>(41/63) | FFPE | IHC IL17<sup>+</sup> cells | ≥ mean value (data not<br>shown) | 39 (2-76) | 6 |
| **Pancreatic Cancer (n = 1)** |  |  |  |  |  |  |  |  |  |
| He SB [63] | 2011 | China | Pro. | 46 (31/15) | FFPE | IHC IL17<sup>+</sup> cells | ≥ median value(not<br>shown) | 5-48 | 8 |
| **CRC (n = 5)** |  |  |  |  |  |  |  |  |  |
| Liu JK [16] | 2011 | China | Pro. | 52 (31/21) | FFPE | IHC IL17<sup>+</sup> cells | ≥ median value (data<br>not shown) | NR | 6 |
| Li XY [20] | 2015 | China | Retro. | 95 (52/43) | FFPE | IHC IL17A<sup>+</sup> cells | score ≥ 2<sup>5</sup> | NR | 7 |
| Lin Y [44] | 2015 | China | Pro. | 78 (46/32) | FFPE | IHC IL17A<sup>+</sup> cells | score ≥ 3 | NR | 7 |
| Li YX [45] | 2015 | China | Retro. | 40 (24/16) | FFPE | IHC IL17A<sup>+</sup> cells | score ≥ 2 | NR | 7 |
| Chen JX [61] | 2014 | China | Pro. | 102 (NR) | FFPE | IHC IL17<sup>+</sup> cells | ≥ 8.39% (mean value) | NR | 7 |
| **Cervical cancer (n = 1)** |  |  |  |  |  |  |  |  |  |
| Yu Q [55] | 2014 | China | Pro. | 153<br>(0/153) | FFPE | IHC IL17A<sup>+</sup> cells | "minimum p value"<br>approach | NR | 8 |
| **Ovarian cancer (n = 1)** |  |  |  |  |  |  |  |  |  |
| Lan CY [33] | 2013 | China | Retro. | 104<br>(0/104) | FFPE | IHC IL17<sup>+</sup> cells | "minimum p value"<br>approach | NR | 7 |
| **Glioblastoma (n = 1)** |  |  |  |  |  |  |  |  |  |
| Cui XL [64] | 2013 | China | Pro. | 41 (18/23) | FFPE | IHC IL17<sup>+</sup> cells | ≥ 15% | 12.9 (4-24) | 7 |

**Abbreviations:** NOS scale, Newcastle-Ottawa quality assessment scale; HNC, Head and neck cancers; NSCLC, Non-small-cell lung cancer; HCC, Hepatocellular Carcinoma; CRC, Colorectal cancer; pro., Prospective study; retro., Retrospective study; NR, Not reported; FFPE, Formalin-fixed, paraffin-embedded tumor tissue; PBMC, Peripheral blood mononuclear cells; PMA, phorbol 12-myristate 13-acetate; IHC, Immunohistochemistry; FC, Flow cytometry.
<sup>1</sup>scoring of immunoreactive intensity: 0 = negative; 1 = weak/moderate; 2 = strong; the scoring of positive cells ratio: 0: <10%; 1: 11-30%; 2: 31-70%; 3: > 71%. The product of two scores ≥ 1 was classified as high IL-17 expression. <sup>2</sup>scoring of immunoreactive intensity: 0 = negative; 1 = weak; 2 = moderate; 3 = strong; 4 = very strong; and scoring of positive cells ratio: 0: < 5%; 1: 6-25%; 2: 26-50%; 3: 51-75%; 4: > 75%. The product of two scores > 6 was classified as high IL-17 expression. <sup>3</sup>The area of staining was scored: 0:<10%; 1: 10–33%; 2: 34–66% and 3:>66%. Cases with a score ≥2 were classified as positive. <sup>4</sup>The “minimum P value” approach, calculated by X-tile software (Yale University, New Haven, CT), was used to assess the cutoff for the best separation of immune markers referring to outcome of patients. <sup>5</sup>same score grading system with <sup>2</sup>, while the product of two scores ≥ 2 was classified as high IL-17 expression.

**Table 2.** Main characteristics of the included studies about Th17 cells in this meta-analysis.

| First author (ref) | Year | Country | Study design | Number (M/F) | Sample | Measurement | Cut-off value (Th17 cells high) | Median follow-up (m) | NOS scale |
| --- | --- | --- | --- | --- | --- | --- | --- | --- | --- |
| **HNC (n = 1)** |  |  |  |  |  |  |  |  |  |
| Lee MH [56] | 2018 | China | Pro. | 120 (99/21) | PBMC + PMA + ionomycin + GolgiPlug 4h | FC CD4<sup>+</sup>IL17A<sup>+</sup> cells | ≥ 3.015% (median value) | 60 | 9 |
| **NSCLC (n = 2)** |  |  |  |  |  |  |  |  |  |
| Song L [18] | 2019 | China | Pro. | 66 (NR) | PBMC + PMA + ionomycin + brefeldin A 4h | FC CD4<sup>+</sup>IL17<sup>+</sup> cells | ≥ 1.15% (median value) | 13.5 (1-87) | 9 |
| Huang J [65] | 2017 | China | Pro. | 54 (37/17) | fresh tumor tissues | FC CD3<sup>+</sup>CD4<sup>+</sup>IL-17<sup>+</sup> cells | ≥ 1.86% (median value) | NR | 7 |
| **Cancers of digestive system (n = 4)** |  |  |  |  |  |  |  |  |  |
| **HCC (n = 2)** |  |  |  |  |  |  |  |  |  |
| Yan J [19] | 2014 | China | Pro. | 150 (NR) | PBMCs + leukocyte activation cocktail 5h | FC CD4<sup>+</sup>IL17<sup>+</sup> cells | ≥ mean value (data not shown) | NR | 8 |
| Yang XW [66] | 2019 | China | Pro. | 26 (15/11) | Malignant ascites + PMA + ionomycin + GolgiStop 5h | FC CD4<sup>+</sup>IL17<sup>+</sup> cells | ≥ 2.75% (median value) | NR | 8 |
| **CRC (n = 1)** |  |  |  |  |  |  |  |  |  |
| Limagne E [67] | 2016 | France | Pro. | 25 (NR) | PBMC + PMA + ionomycin + monensin | FC CCR6<sup>+</sup>CXCR3<sup>-</sup> cells | ≥ mean value (data not shown) | NR | 7 |
| **Gallbladder cancer (n = 1)** |  |  |  |  |  |  |  |  |  |
| Patil RS [68] | 2016 | India | Pro. | 40 (NR) | PBMC + PMA + ionomycin 5h | FC CD4<sup>+</sup>IL17A<sup>+</sup> cells | ≥ 1.80% (mean value) | NR | 7 |
| **Leukemia (n = 3)** |  |  |  |  |  |  |  |  |  |
| Abousamra NK [69] | 2013 | Egypt | Pro. | 93 (51/42) | PBMC + PMA + ionomycin + BFA 5h | FC CD3<sup>+</sup>CD4<sup>+</sup>IL-17<sup>+</sup> cells | ≥ 2.90% (median value) | NR | 8 |
| Jain P [70] | 2011 | America | Pro. | 55 (NR) | PBMC + PMA + ionomycin + monensin 5h | FC CD3<sup>+</sup>CD4<sup>+</sup>IL-17<sup>+</sup> cells | ≥ 5.80% (median value) | NR | 7 |
| Han YX [71] | 2014 | China | Pro. | 98 (NR) | BMMCs + PMA + ionomycin + brefeldin A 5h | FC CD3<sup>+</sup>CD4<sup>+</sup>IL-17<sup>+</sup> cells | ≥ 3.41% (median value) | NR | 7 |

**Abbreviations:** PBMC, Peripheral blood mononuclear cells; PMA, phorbol
12-myristate 13-acetate; BMNCs, bone marrow mononuclear cells.

**Table 3.** Summary of the clinical outcomes in the meta-analysis.

| Group | No. of studies | No. of patients | RR/HR[95% CI] | P for heterogeneity | I<sup>2</sup> | References |
| --- | --- | --- | --- | --- | --- | --- |
| **IL-17<sup>+</sup> cells[74-85]** |  |  |  |  |  |  |
| Histological grade (poor vs. well/moderate) | 14 | 1645 | 1.10 [0.90, 1.34] | 0.01 | 53.3% | [16,20,28,42-47] |
| Lymph node metastasis | 9 | 1074 | 0.97 [0.81, 1.15] | 0.038 | 50.9% | [17,20,38,42-46,48] |
| Tumor volume<br>(T3/4 vs. T1/2) | 8 | 1036 | 1.04[0.85, 1.26] | 0.003 | 67.2% | [17,20,38,42,43,46,48,49] |
| Clinical stages<br>(III/IV vs. I/II) | 11 (Total) | 1370 | 1.30 [0.99, 1.70] | <0.001 | 73.9% | [17,20,40-43,45,46,48-50] |
|  | NSCLC (n = 2) | 158 | 2.34 [1.30, 4.18] | 0.994 | 0.0% | [40,41] |
| Distant metastasis | 6 | 602 | 1.36 [0.70, 2.63] | 0.016 | 64.0% | [38,39,42,44,46,49] |
| Tumor recurrence | 4 | 317 | 4.23 [1.58, 11.35] | 0.017 | 70.5% | [38,39,44,49] |
| DFS | 9 (Total) | 1448 | 1.84 [1.22, 2.77] | <0.001 | 87.7% | [17,19,40,46,51-55] |
|  | Cancers of digestive system (n = 6) | 1032 | 2.23 [1.65, 3.01] | 0.005 | 70.5% | [19,46,51-54] |
| OS | 25 (Total) | 3439 | 1.39 [1.04, 1.87]) | <0.001 | 97.0% | [40-43,49,50,53,54,56-62] |
|  | HNC (n = 2) | 200 | 2.98 [1.59, 5.57] | 0.421 | 0.0% | [40-43,49,50,53,54,56-62] |
|  | Cancers of digestive system (n = 18) | 2851 | 1.45 [1.03, 2.04] | <0.001 | 95.5% | [40-43,49,50,53,54,56-62] |
| **Th17 cells** |  |  |  |  |  |  |
| OS | 10 | 727 | 0.93 [0.58, 1.49] | <0.001 | 86.1% | [18,19,56,63-69] |

### IL-17^+^ cells and histological grade

As shown in Fig 2.A, the relevance between high level of IL-17^+^ cells and histological grade (poor differentiation versus (vs.) well/moderate differentiation) was evaluated in 14 studies including head and neck cancers (HNC) (n = 2) [38,39], breast cancer (n = 1) [17], non-small-cell lung cancer (NSCLC) (n = 2) [40,41] and cancers of digestive system (n = 9) [16,20,28,42–47], with 1645 patients. Results showed no statistical significance was found between high level of IL-17^+^ cells and histological grade (RR = 1.10, 95% CI [0.90, 1.34]). Due to the heterogeneity, a random effect model was applied (*I*^2^ = 53.3%, p = 0.01).

**Fig 2.**
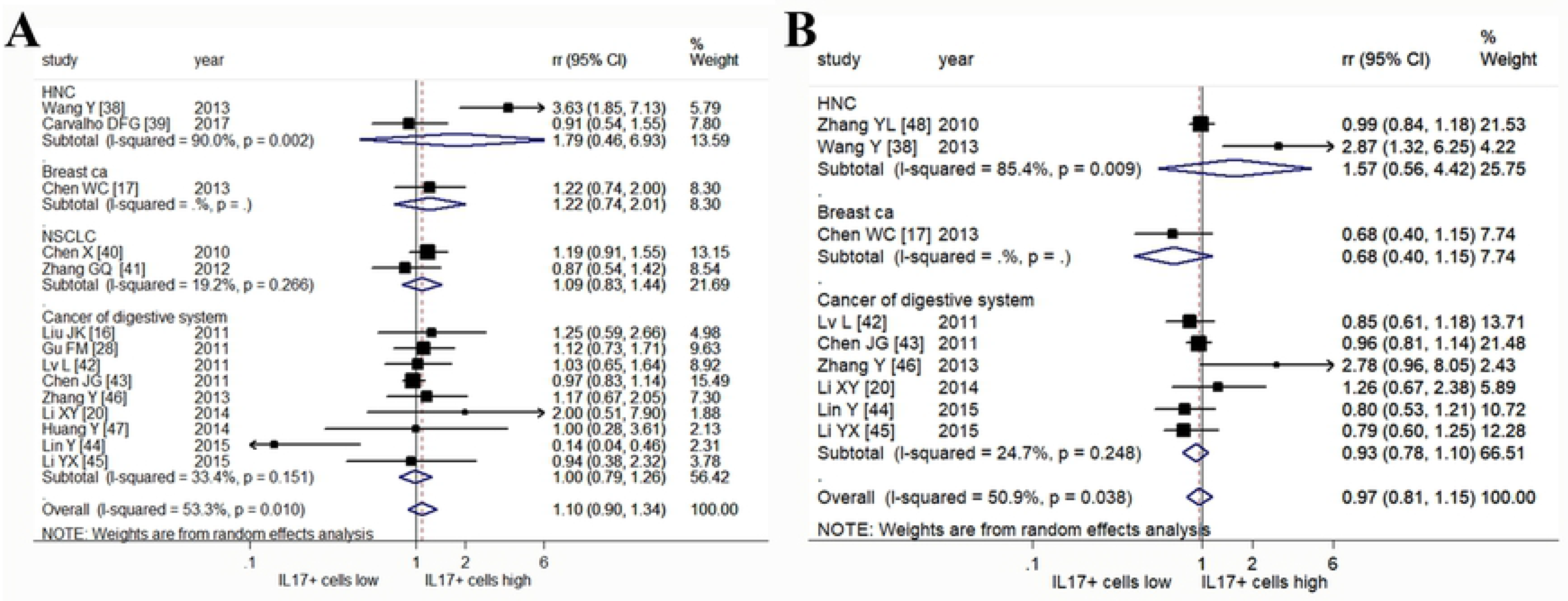
Forrest plots of risk ratios (RRs) for correlation between high IL-17^+^ cells and clinicopathological features. (A) Histological grade and (B) lymph node metastasis.

### IL-17^+^ cells and lymph node metastasis

The association between high IL-17^+^ cells and lymph node metastasis was explored in 9 studies with 1074 patients [17,20,38,42–46,48] (Fig 2.B). RR for the relevant 9 studies was 0.97 (95% CI [0.81, 1.15]), indicating the high IL-17^+^ cells within tumor tissues had no significant influence on lymph node status. Considering the heterogeneity, a random effect model was used (*I*^2^ = 50.9%, p = 0.038).

### Prognostic values of IL-17^+^ cells in tumor volume and clinical stages

The prognostic values of IL-17+ cells in tumor volume (T3/T4 vs. T1/T2 stages)(Fig 3.A) and clinical stage (III/IV vs. I/II stages)(Fig 3.B) were investigated in 8 studies with 1036 patients [17,20,38,42,43,46,48,49] and 11 studies with 1370 patients [17,20,40–43,45,46,48–50], respectively. Pooled RR indicated there was no significant correlation between high IL-17^+^ cells and tumor volume (RR = 1.04, 95% CI [0.85, 1.26]). The high IL-17^+^ cells was apt to cause advanced clinical stages, with a pooled RR of 1.30, though a statistical significance was not appeared (95% CI [0.99, 1.70]). Besides, result of subgroup analysis showed high level of IL-17^+^ cells in NSCLC was contributed to advanced clinical stages in 2 studies with 158 patients (RR = 2.34, 95% CI [1.30, 4.18]) [40,41]. The random effect model was applied for analyses of tumor volume and clinical stages owing to the obvious heterogeneities (*I*^2^ = 67.2%, p = 0.003 and *I*^2^ = 73.9%, p < 0.001, respectively).

**Fig 3.**
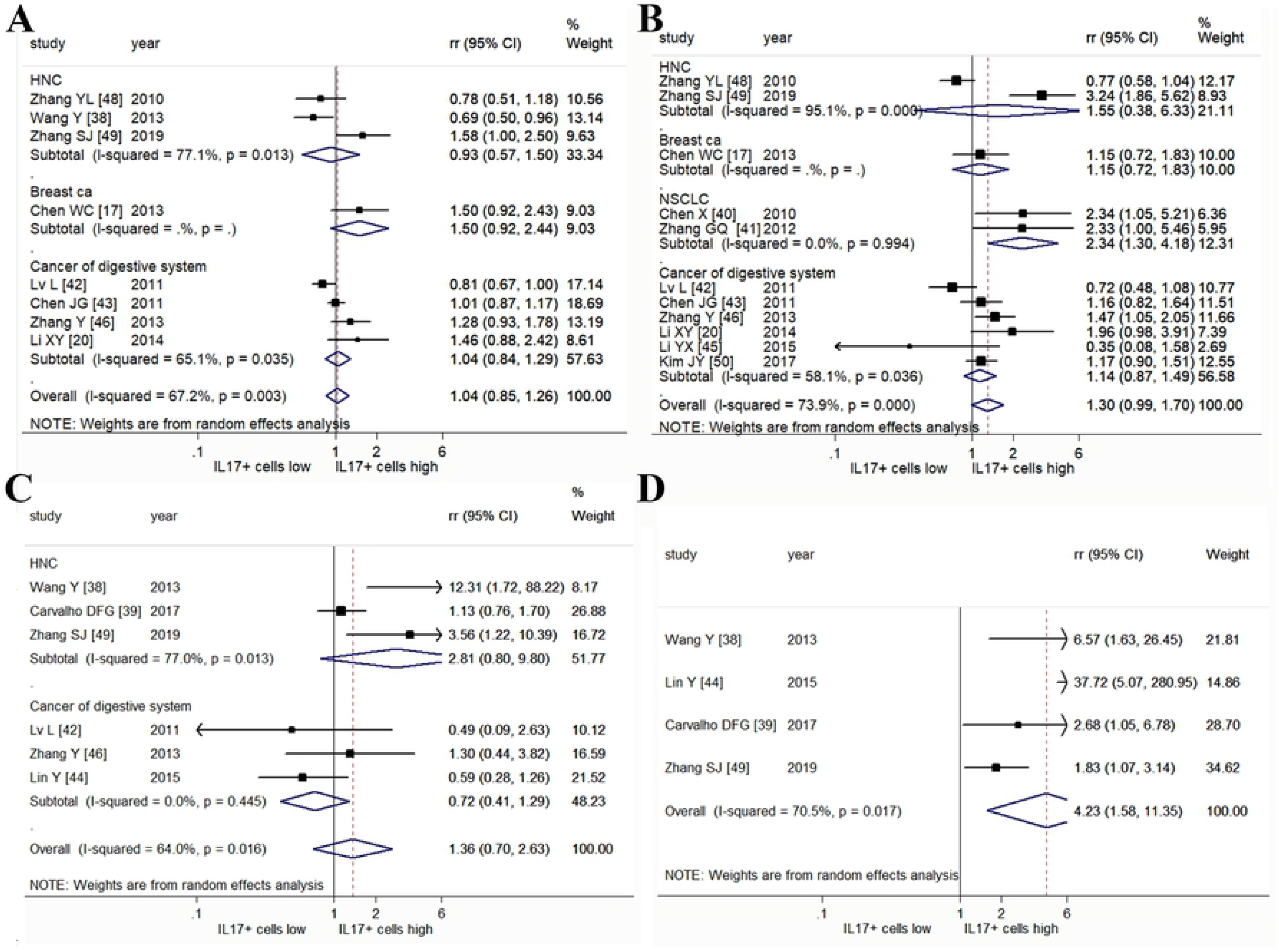
Forrest plots of RRs for correlation between high IL-17^+^ cells and clinicopathological features. (A) Tumor volume (T stages), (B) clinical stages, (C) distant metastasis and (D) tumor recurrence.

### Distant metastasis and tumor recurrence

We further analyzed the impact of IL-17^+^ cells on distant metastasis (Fig 3.C) and tumor recurrence (Fig 3.D). Pooled RR of 6 studies with 602 patients showed a non-statistically significant correlation between high IL-17^+^ cells and distant metastasis (RR = 1.36, 95%CI [0.70, 2.63]) [38,39,42,44,46,49]. Moreover, 4 studies with 317 patients investigated the correlation between high IL-17^+^ cells and tumor recurrence [38,39,44,49]. Results indicated the high IL-17^+^ cells was significantly related to tumor recurrence (RR = 4.23, 95% CI [1.58, 11.35]). Considering the obvious heterogeneities, we used the random effect model in analysis of distant metastasis (*I*^2^ = 64.0%, p = 0.016) and tumor recurrence (*I*^2^ = 70.50%, p = 0.017).

### IL-17^+^ cells and DFS

As shown in Fig 4.A, high IL-17^+^ cells within tumor tissues was proved to be an unfavorable indicator of DFS in 9 studies with 1448 patients (HR = 1.84, 95% CI [1.22, 2.77]) [17,19,40,46,51–55]. In subgroup analysis, high IL-17^+^ cells within cancers of digestive system was significantly connected with worse DFS, with a pooled HR of 2.23 (95% CI [1.65, 3.01]) in 6 studies with 1032 patients [19,46,51–54]. The random effect model was used because of significant heterogeneity (*I*^2^ = 87.7%, p < 0.001).

**Fig 4.**
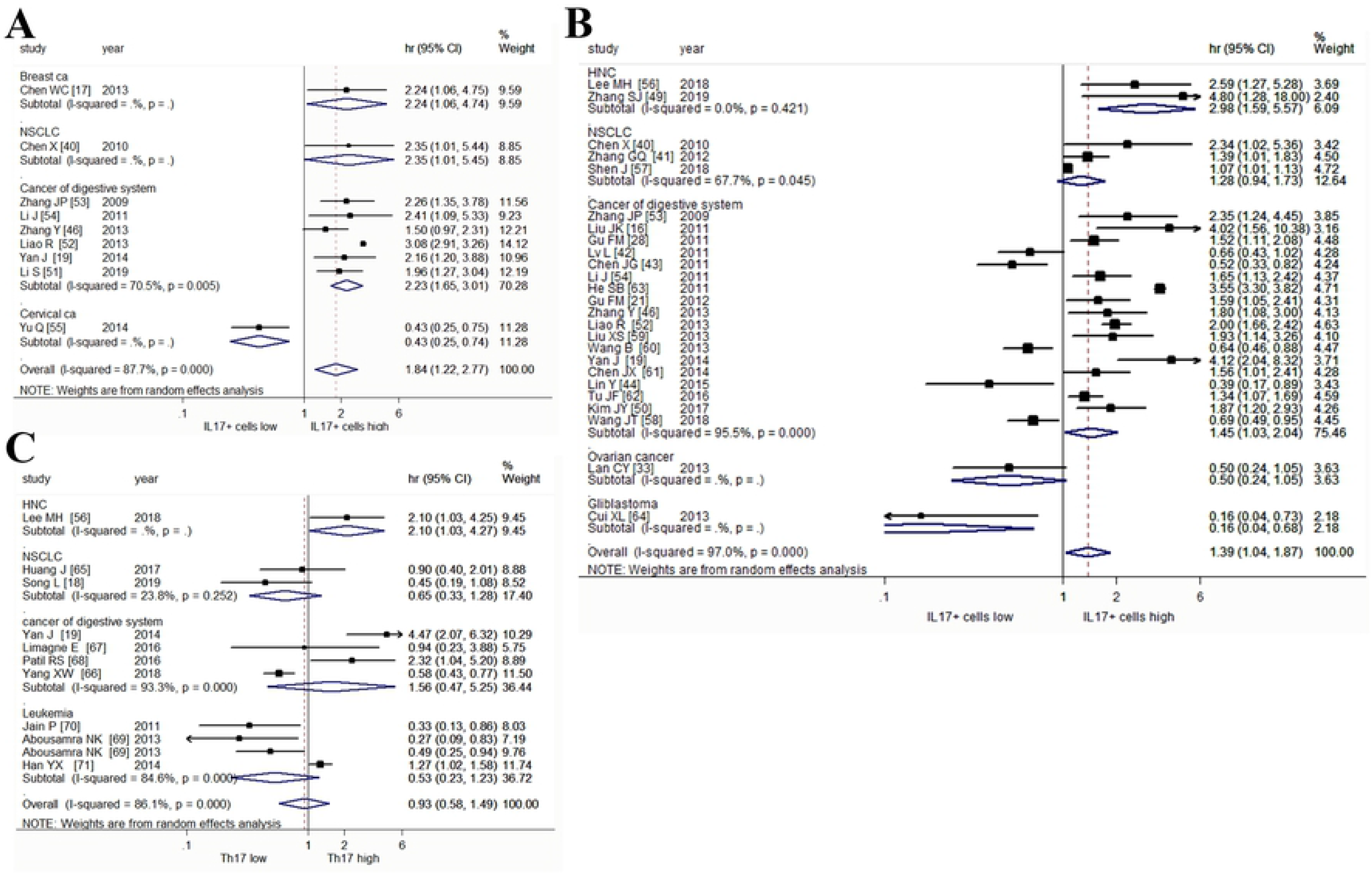
Forrest plots of hazard ratios (HRs) for high IL-17^+^ cells about survival outcomes of (A) disease free survival (DFS) and (B) overall survival (OS). (C) HR correlation between high Th17 cells OS.

### IL-17^+^ cells and OS

Collectively, the pooled HR of 1.39 (95% CI [1.04, 1.87]) which aggregated from 25 studies with 3439 patients indicated high IL-17^+^ cells within tumor tissues was associated with significantly poorer OS (Fig 4.B) [40–43,49,50,53,54,56–62]. The results of subgroup analysis showed high IL-17^+^ cells within tumor tissues was an unfavorable marker of OS in both HNC (HR = 2.98, 95% CI [1.59, 5.57]) (n = 2, with 200 patients) and cancers of digestive system (HR = 1.45, 95% CI [1.03, 2.04]) (n = 18, with 2851 patients). The random effect model was performed in consideration of the obvious heterogeneity (*I*^2^ = 97.0%, p < 0.001).

### Th17 cells and survival

Finally, we also investigated the impact of Th17 cells on OS in 10 studies with 727 patients (Fig 4.C) [18,19,56,63–69]. The Th17 cells were identified by flow cytometery analysis in all included studies. Most of included studies analysis Th17 cells were sampled from peripheral blood mononuclear cells (PBMC), of which one study sampled from fresh tumor tissues [63] and another study sampled from malignant ascites [64]. Pooled HR showed no statistically significant association was discovered between high Th17 cells and OS (HR=0.93, 95% CI [0.58, 1.49]). A random effect model was performed in consideration of the obvious heterogeneity (*I*^2^ = 86.1%, p < 0.001).

### Publication bias

The statistics about publication bias were evaluated by Begg’s funnel plot and Egger’s test (Fig 5). No publication bias was observed by in Begg’s funnel plots of histological grade (p = 0.702), lymoh node metastasis (p = 0.532), tumor volume (p = 0.322), clinical stages (p = 0.312), distant metastasis (p = 0.573), DFS (p =0.297), OS of both IL-17^+^ cells (p = 0.102) and Th17 cells (p = 0.186). Moreover, results of Egger’s test also demonstrated no publication bias was founded in histological grade (p = 0.757), lymph node metastasis (p = 0.358), tumor volume (p = 0.345), clinical stages (p = 0.297), distant metastasis (p = 0.366), OS of both IL-17^+^ cells (p = 0.655) and Th17 cells (p = 0.724). The potential publication bias was observed in tumor recurrence (Begg’s funnel plot of p = 0.042 and Egger’s test of p = 0.031) and DFS of IL-17^+^ cells with a p value of 0.039 in Egger’s test.

**Fig 5.**
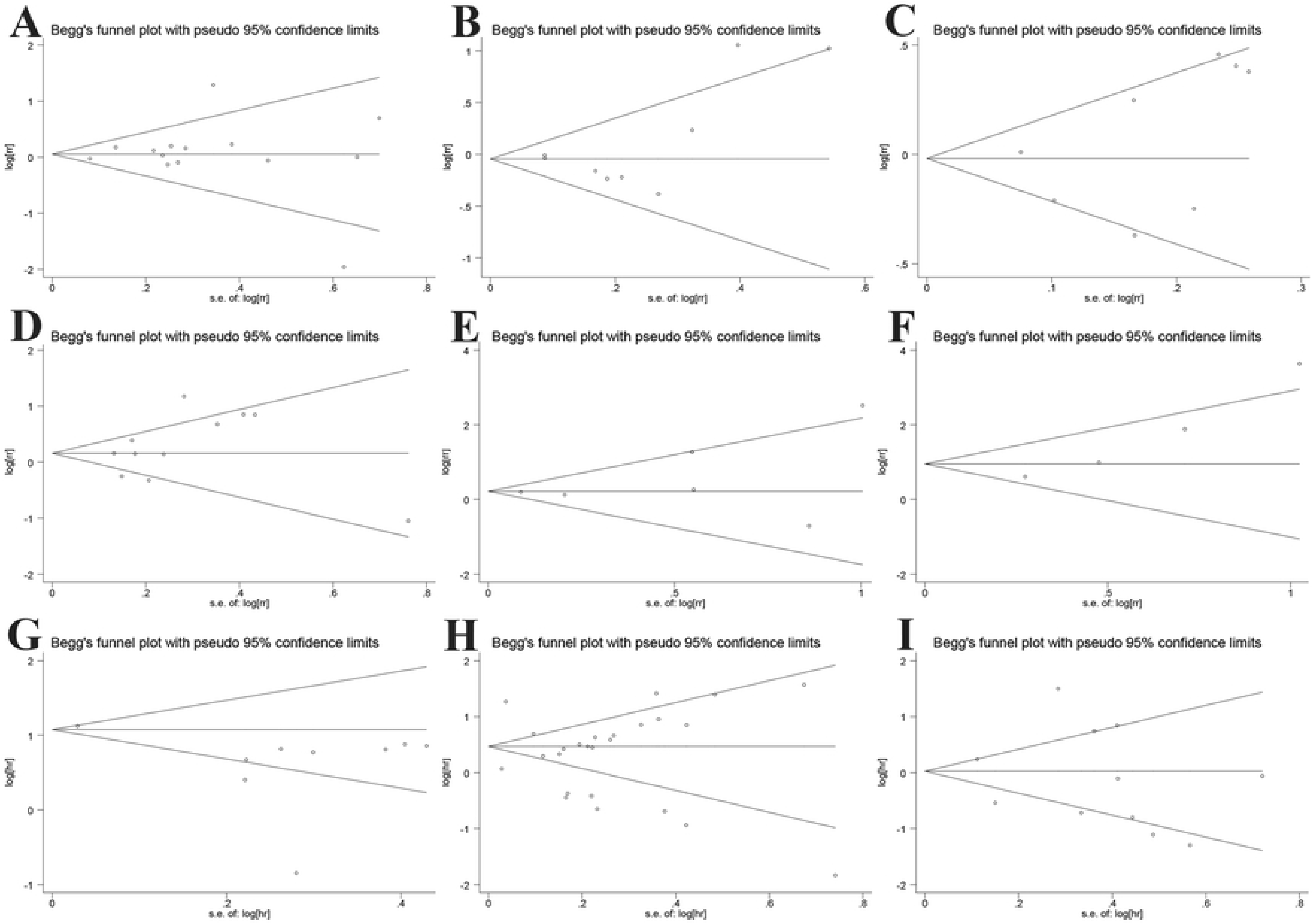
Funnel graph for evaluating the potential publication bias in this meta-analysis. No potential publication bias was observed except analyses of tumor recurrence and DFS. (A) Histological grade, (B) lymph node metastasis, (C) tumor volume (T stages), (D) clinical stages, (E) distant metastasis, (F) tumor recurrence, (G) DFS about IL-17^+^ cells, (H) OS about IL-17^+^ cells and (I) OS about Th17 cells.

## Discussion

The Th17 or IL-17 dominance is usually related to autoimmune diseases such as psoriasis, ankylosing spondylitis (AS), rheumatoid arthritis (RA) or Systemic Lupus Erythematosus (SLE) [70]. The role of Th17 or other IL-17^+^ cells in cancer progression is often linked to inflammation that mediated by themselves. Th17 cells or other IL-17^+^ cells can play a dual role in both promoting tumor development and tumor suppression, though the underlying mechanism still largely remains elusive. A considerable number of clinical studies have discussed the correlations between IL-17^+^/Th17 cells and outcomes of cancer patients, while the conclusions are still controversial. Therefore, this current study was carried out for a deeper understanding of values of Th17 cells and IL-17^+^ cells in the prognosis of cancer patients.

To our knowledge, it is the first comprehensive and most full-scale meta-analysis to demonstrate the underlying values of elevated IL-17^+^ cells and Th17 cells levels in human cancers. In summary, high IL-17^+^cells was shown to be significantly correlated with tumor recurrence, worse DFS and OS, while no significant difference was observed between high IL-17^+^ cells and histological grade, lymph node metastasis, tumor volume, clinical stages or distant metastasis. Notably, results of subgroup analysis further suggested high IL-17^+^ cells was significantly associated with advanced clinical stages in NSCLC. Moreover, pooled HRs demonstrated that high IL-17^+^ cells was an unfavorable indicator of DFS and OS especially in cancers of digestive system, and also represented worse OS in HNC. Furthermore, we could only accomplished the meta-analysis about Th17 cells level and OS owing to the limited number of studies about relationships between high Th17 cells and relevant clinicopathologic features. No significant association was observed between high Th17 cells and OS. Among the included studies about IL-17^+^ cells, most of these studies identified IL-17^+^ cells within tumor tissues by immunohistochemistry staining (IHC), while only one study identified IL-17^+^ cells from PBMC by flow cytometry (FC) [56]. As for Th17 cells, most of the included studies sampled from PBMC, though one study sampled from malignant ascites [64], one from fresh tumor tissues [63] and another one from bone marrow mononuclear cells (BMMCs) [69]. The level of Th17 cells in peripheral blood of cancer patients is frequently founded to be higher than that in normal control, which might suggest an underlying inflammatory status [35]. Interesting, the high Th17 cells sampled from peripheral blood of patients with leukemia was correlated with better survival [67,68], while high Th17 cells sampled from bone marrow of patients with leukemia was an indicator of worse survival [69]. Thus, it might not properly reflect the actual role of Th17 cells in tumor microenvironment purely by investigating the role Th17 cell in peripheral blood. Besides, the increased accumulation of Th17 cells in malignant ascites was reported to indicate improved patient survival [64]. Overall, the actual role and underlying mechanism of Th17 cells in tumor microenvironment still remain largely unclear, while IL-17^+^ cells seem to play a pro-tumor function. Therefore, it will be worthwhile to further investigate the potential role of primary Th17 cells in tumor microenvironment.

A tumor-promoting function of IL-17 expression has been observed in inflammatory related and sporadic cancers of gastric, liver and colon [71]. Increased IL-17 expression has been shown to be associated with resistance to chemotherapy and target therapy, such as anti-VEGF drug (Avastin) [12,72]. The anti-VEGF treatment induced the accumulations of Th17 cells and other IL-17^+^ cells in tumor microenvironment, of which the elevated IL-17 level facilitated the recruitment of myeloid-derived suppressor cells (MDSCs) with immune-suppressive and proangiogenic functions, thus leading resistance to anti-VEGF target therapy in lymphoma, lung and colon cancers [72]. Wang et al. have reported the treatment of fluorouracil (5-FU) for mice with CRC caused an increased IL-17A expression within tumors, which might weak the response to chemotherapy. Importantly, the co-administration of IL-17A antibody, which by itself was insufficient to inhibit tumor, was able to generate synergetic effect with 5-FU treatment to enhance therapeutic response [12]. Consistent with the conclusion in our meta-analysis, patients with cancers of digestive system whose tumors display with high IL-17^+^ cells are associated with poorer prognosis and shorter OS [73]. These patients are the ones who most likely to be benefit from the target therapy of IL-17 blockade. However, more relevant clinical evidences are urgently needed before any conclusion can be drawn.

Some limitations still could not be ignored in this meta-analysis. First, although all of the included studies used IHC for IL-17^+^ cells and flow cytometry for Th17 cells to assess their expression levels, the diverse cut-off values might be the potential reasons to cause publication bias. Nevertheless, subgroup analysis according to different cut-off values is unable to implement, due to the limited number of corresponding studies. Besides, we carried out subgroup analysis according to various cancer types in order to reduce potential publication bias and make results more reliable.

Furthermore, obvious heterogeneities were also observed in this meta-analysis, which might be generated by many causes such as different cancer types, cut-off values, primary antibody sources, follow-up time and sample sizes. Therefore, we further applied the random effect model for decreasing the impact of heterogeneity on outcomes.

In conclusion, our meta-analysis suggests high IL-17^+^ cells could be an indicator for worse survival in patients with malignant cancers, especially in cancers of digestive system. Although the level of Th17 cells appears to have non-statistically significant impact on prognosis of patient with malignant cancers, more clinical researches should be implemented to further investigate the underlying function of Th17 cells within tumor microenvironment. This study put forward a new insight for the potential application of anti-IL-17 target therapy in cancer patients.

## Funding

This work was funded by grant from National Natural Science Foundation of China (81201788).

## Author contributions

Y Luo and T Yu contributed to the data extraction, analysis and manuscript writing. C Yi and HS Shi contributed to the design of study, revising of manuscript and financial support.

---

**S1 Table.** Search strategies in databases.

**S2 Table.** PRISMA Checklist.

## Notes

**Disclosure of Potential Conflict of Interest:** No potential conflicts of interest were disclosed.

